# A serotonergic mechanism is involved in the pro-cognitive effect of AM6545 treatment in a mouse model of fragile X syndrome

**DOI:** 10.1101/2025.01.24.634730

**Authors:** Lucía de los Reyes-Ramírez, Araceli Bergadà-Martínez, Sara Martínez-Torres, Irene Martínez-Gallego, Pilar Losada-Ruiz, Silvia Arboleya, Miguel Gueimonde, Rafael Maldonado, Antonio Rodríguez-Moreno, Andrés Ozaita

## Abstract

Fragile X syndrome (FXS) is the principal monogenic syndrome leading to inherited intellectual disability and autism. It is caused by the silencing of *FMR1* gene that leads to the loss in the expression of its encoded protein, the fragile X messenger ribonucleoprotein 1 (FMRP). In synapses, FMRP has a key role in local mRNA modulation to maintain synaptic plasticity. The *Fmr1* KO (FX) mouse model shows cognitive impairment and some of the synaptic traits present in individuals with FXS together with alterations in gut microbiota. Previous studies revealed that pharmacological and genetic cannabinoid type-1 receptor (CB1R) inhibition significantly prevented central key alterations in FX mice. Here, we aimed to evaluate the effect of a sub-chronic treatment with the largely peripherally-restricted CB1R antagonist AM6545. We found that AM6545 reduced memory deficits and restored enhanced hippocampal mGluR5-dependent long-term depression and aberrant dendritic spine density in FX mice. At the peripheral level, AM6545 modified altered FX mice fecal microbiota composition, while in the hippocampus AM6545 treatment upregulated hippocampal *Htr4*, the gene encoding for serotonin receptor 4 (5-HT4R) This upregulation positively correlated with memory performance. Notably, acute pharmacological blockade of 5-HT4R abolished the pro-cognitive effect produced by AM6545. Together, our results suggest that peripheral CB1R inhibition ameliorates key alterations in FX mouse model and modifies the expression of serotonergic receptors important for cognitive performance.

## Introduction

Fragile X syndrome (FXS) is the most common monogenic origin of inherited intellectual disability and autism spectrum disorder (ASD) [1]. It is caused by a trinucleotide CGG expansion in the 5’ UTR of the *FMR1* gene, which leads to the loss of expression of the fragile X messenger ribonucleoprotein 1 (FMRP) [2]. People with FXS present heterogeneous symptoms, including memory impairment, hyperactivity, impulsivity or autistic-like behavior [3].

The *Fmr1* gene knockout (FX) mouse model recapitulates the main traits of the disorder [4]. Notably, an uncontrolled activity of the metabotropic receptor mGluR5, revealed by an enhanced mGluR5-dependent long-term depression (LTD) has been described in FX mice [5,6], as well as alterations in dendritic spine density and morphology with an increase in the immature dendritic spines [7,8]. In addition to these neuronal traits, the FX mice show gut microbiota dysbiosis [9].

The endocannabinoid system is a key modulator of synaptic plasticity and cognitive function [10], two aspects that are affected in FXS. It was previously reported that systemic cannabinoid type-1 receptor (CB1R) inhibition using genetic or pharmacological approaches ameliorated cognitive deficits in FX mice, as well as the aberrant mGluR5-LTD and the altered CA1 spine density in the hippocampus [7,11]. However, systemic CB1R inhibition was found to induce psychiatric side effects when used as an anti-obesity treatment [12]. To avoid these hypothetical central adverse effects of CB1R inhibition, diverse strategies have been developed including CB1R peripherally-restricted antagonists and allosteric modulators. We recently demonstrated that AM6545, a peripherally-restricted CB1R antagonist, enhances memory performance in naïve mice [13,14]. Therefore, we aimed to explore the potential beneficial effects of AM6545 in the alleviation of key alterations of FX mice.

The present study shows that AM6545 treatment ameliorates object-recognition memory and improves cellular and structural alterations in FX mice. Furthermore, we described a decrease in *Akkermansia* and *Lactobacillus* genera in FX mice that increase after AM6545 treatment. Interestingly, we found that the upregulation of serotonin receptor 4 in FX mice after the treatment is needed for AM6545 mnemonic effect. Together, this evidence undisclosed the inhibition of peripheral CB1R as an approach to alleviate the cognitive deficits in a mouse model for FXS through peripheral noradrenergic and serotonergic mechanisms.

## Materials and methods

### Ethics

Animal procedures were conducted following “Animals in Research: Reporting Experiments” (ARRIVE) guidelines and standard ethical guidelines [15,16] (European Directive 2010/63/EU), and were approved by the local ethical committee (Comitè Ètic d’Experimentació Animal-Parc de Recerca Biomèdica de Barcelona, CEEA-PRBB).

### Animals

FX mice in Friend Virus B background (*Fmr1* KO, FVB.129P2-Pde6b+Tyrc-chFmr1tm1Cgr/J) and wild-type mice (WT, FVB.129P2-Pde6b+Tyrc-ch/AntJ) were purchased from The Jackson Laboratory and crossed to obtain outbred FX and WT littermates. FX mice in C57BL/6J congenic background (B6.129P2-Fmr1tm1Cgr/J) were obtained from the Baylor College of Medicine Mouse Facility and crossed to generate inbred FX and WT littermates. The transgenic line Tg(Thy1-EGFP) MJrs/J (Stock #007788, The Jackson Laboratories, USA) crossed to FX mice in C57BL/6J background was used to study the density and morphology of dendritic spines. Young-adult male mice (12-14 weeks) used in this study were housed in Plexiglas cages (2-4 mice/cage) and maintained in temperature (21 ± 1°C) humidity (55 ± 10%) and light (light on at 8 am; light off at 8 pm) controlled environment. Food and water were available *ad libitum*. Experiments were performed during the light phase after animals were handled for 1 week and habituated to the experimental room. Behavioral experiments were conducted by an observer blind to experimental conditions.

### Drugs and treatments

AM6545 (1 mg/kg) from Tocris-Bio-Techne was dissolved in 0.26% DMSO, 4.74% ethanol, 5% cremophor-EL and 90% saline. Sotalol (10 mg/kg) from Sigma-Aldrich was dissolved in saline [13]. GR125487 (10 mg/kg) from Tocris-Bio-Techne was dissolved in saline. Drugs were injected intraperitoneally (i.p.) in a volume of 10 ml/kg of body weight for acute or sub-chronic (7d) treatment.

### Novel object-recognition test

NORT was performed as previously described [13]. For sub-chronic treatment, the last AM6545 administration was performed after the familiarization phase (Supplementary Fig. 1A). Sotalol was administered acutely after the familiarization phase and, when required, 20 min before AM6545 acute administration (Supplementary Fig. 1B). GR125487 was administered acutely after the familiarization phase and, when required, 20 min before the last AM6545 administration (Supplementary Fig. 1C).

### Slice preparation for electrophysiological recordings

Twenty-four hours after the last administration of AM6545 (1 mg/kg, 7d, i.p.), WT and FX mice were anesthetized with isoflurane (5%) and decapitated for slice preparation. Hippocampal slices were prepared as previously described [17–19].

### Electrophysiological recordings

Whole-cell patch-clamp recordings were made from pyramidal cells located in the CA1 field of the hippocampus. Cells were patched under visual guidance by infrared differential interference contrast microscopy and verified to be pyramidal neurons by their characteristic voltage response to a current step protocol. Neurons were recorded in voltage-clamp configuration with a patchclamp amplifier (Multiclamp 700B), and data were acquired using pCLAMP 10.2 software (Molecular Devices). Patch electrodes were pulled from borosilicate glass tubing and had a resistance of 4–7 MΩ when filled with (in mM) CsCl, 120; HEPES, 10; NaCl, 8; MgCl2, 1; CaCl2, 0.2; EGTA, 2; and QX-314, 20 (290 mOs/ml, pH 7.2–7.3). Cells were excluded from analysis if the series resistance changed by more than 15% during the recording. Recordings were low-pass filtered at 3 kHz and acquired at 10 kHz. mGluR5-LTD was induced by treating slices with 3,5-dihydroxyphenylglycerine (DHPG) (100 μM) for 10 min. LTD was quantified by comparing the mean EPSC amplitude over the 30 min post-drug period with the mean EPSC amplitude during the baseline period and calculating the percentage change from 5 last min.

### Brain slice preparation for dendritic spine analysis

Twenty-four hours after the last administration of AM6545 (1 mg/kg, 7d, i.p.), mice were deeply anesthetized by i.p. injection (0.2 ml/10 g of body weight) of a mixture of ketamine/xylazine (100 mg/kg and 20 mg/kg, respectively) prior to intracardiac perfusion of cold 4% paraformaldehyde. Brains were removed and stored in a solution of 30% sucrose at 4°C. Coronal frozen sections were made at 60 μm on a freezing microtome (Leica) and mounted for image acquisition.

### Dendritic spine analysis

Secondary and tertiary apical dendrites of pyramidal neurons from the *stratum radiatum* of CA1 region of the dorsal hippocampus were analyzed. Images were acquired with a confocal microscope (TCS SP5 STED Leica) using a glycerol immersion lens plus 3 times magnification (63X/3) with a sequential line scan at 1024 × 1024 pixel resolution. Serial optical sections were acquired with a 0.13 μm step size. A minimum of 8 dendrites per animal were selected from 4 different slices. A maximum of 2 dendrites were selected from the same neuron. Confocal images were deconvoluted using Huygens Essential software and dendritic spines were analyzed and classified using the semiautomated software NeuronStudio with the described parameters (Supplementary Table 1). Spine density was calculated by expressing the average number of spines in a 10 μm portion of the dendrite. Each spine was checked manually by an observer blind to experimental conditions for unbiased classification

### Sample preparation and 16S rRNA gene profiling of fecal microbiota analysis

Fresh fecal samples were collected from each experimental group (Supplementary Table 2), and stored at −80°C until used. Fecal pellets were subsequently diluted 1/10 (w/v) in phosphate buffer saline and vigorously homogenized by vortexing. Cell pellet were isolated by centrifugation at 10,000 rpm for 15 min and used for bacterial DNA isolation using QIAamp DNA stool kit (Qiagen, GmbH, Germany) following manufacturer’s instructions, including a bead-beating step in a Fastprep (MP Biomedicals, Thermo Fisher Scientific Inc.). The isolated DNA was used as template for partial sequencing (hypervariable V3-V4 region) of the *16S* rRNA gene on the Illumina platform following in-house protocols (Novogene Co.) [20]. Data from *16S* profiling gut microbiota were total sum scaling and cumulative sum scaling normalization for diversity and abundance analyses, respectively [21]. α-diversity was studied by Chao 1 and Shannon indexes calculated with *QIIME*.

### Tissue preparation for mRNA analysis

Twenty-four hours after the last AM6545 administration (1 mg/kg, 7d, i.p.), mice were euthanized by cervical dislocation and hippocampal tissues were dissected on ice, frozen on dry ice and stored at −80°C until used.

### Synaptoneurosome preparation for RNA extraction

Synaptoneurosomes enriched fractions were obtained as previously described [22]. RNA extraction was performed using RNeasy Mini kit (74140 QIAGEN) according to the manufacturer’s instructions and RNA concentration and integrity were measured using Agilent 2100 Bioanalyzer (Agilent Technologies).

### RNA-seq and differential gene expression analysis

RNA libraries from hippocampal synaptoneurosomes were obtained and processed by Macrogen as previously described [14]. Raw-data is available at GSE278037. Differential gene expression (DGE) between experimental groups (Supplementary Table 2) was analyzed using *DESeq2* (v1.30) [23] with the cut-off set on |log_2_fold change| > 0.2 and *p-value* < 0.1 after Wald test correction procedure.

### Reverse transcription and quantitative real-time PCR analysis

Reverse transcription and quantitative real-time PCR (qRT-PCR), were performed for hippocampal synaptoneurosomes RNA as previously described [14]. Quantification was performed using the comparative CT Method, relative expression was normalized to *18S* and the fold change was calculated using the 2(–ΔΔCt) formula [24]. The following primers were used: *Htr4*F - TGGAACAACATCGGCATAGTTGA and *Htr4*R - GTGATGGCATAGGGCTTGTTGAC [25], and *18S*F - GTAACCCGTTGAACCCCATT and *18S*R - CCATCCAATCGGTAGTAGCG [26].

### Statistical analysis

Data analyses were performed using GraphPad Prism 8, RStudio (v4.0), MicrobiomeAnalyst and IBM SPSS (v26.0) software. Statistical comparisons were evaluated using unpaired Student’s t-test for two groups comparison, Mann-Whitney U test for two groups comparisons in microbiota experiments (non-normally distributed data) or two-way ANOVA for multiple comparisons. Subsequent Bonferroni *post hoc* was used when required (significant interaction between factors). Results were expressed as mean ± s.e.m. Comparisons were considered statistically significant when *p* < 0.05.

## Results

### AM6545 treatment ameliorate memory deficits in FX mice through a peripheral noradrenergic mechanism

First, we tested whether sub-chronic AM6545 treatment could improve memory deficits in FX mice. WT and FX mice received the last administration of a sub-chronic treatment with AM6545 (1 mg/kg, 7d) immediately after the familiarization phase in the NORT and the test phase was performed 24h later. AM6545 ameliorated cognitive impairment in FX mice both in FVB.129 (two-way ANOVA, interaction: F(1,38) = 4.723, *p* = 0.036; *post hoc* Bonferroni, WT-VEH *vs.* FX-VEH *p* = 0.005; FX-VEH *vs.* FX-AM6545 *p* = 0.001) (Fig. 1A) and C57BL/6J genetic background (two-way ANOVA, interaction: F(1,40) = 10.49, *p* = 0.002; *post hoc* Bonferroni, WT-VEH *vs.* FX-VEH *p* = 0.0004; FX-VEH *vs.* FX-AM6545 *p* = 0.0004) (Fig. 1B), reinforcing the significance of the pro-cognitive effect in both backgrounds.

**Figure 1:**
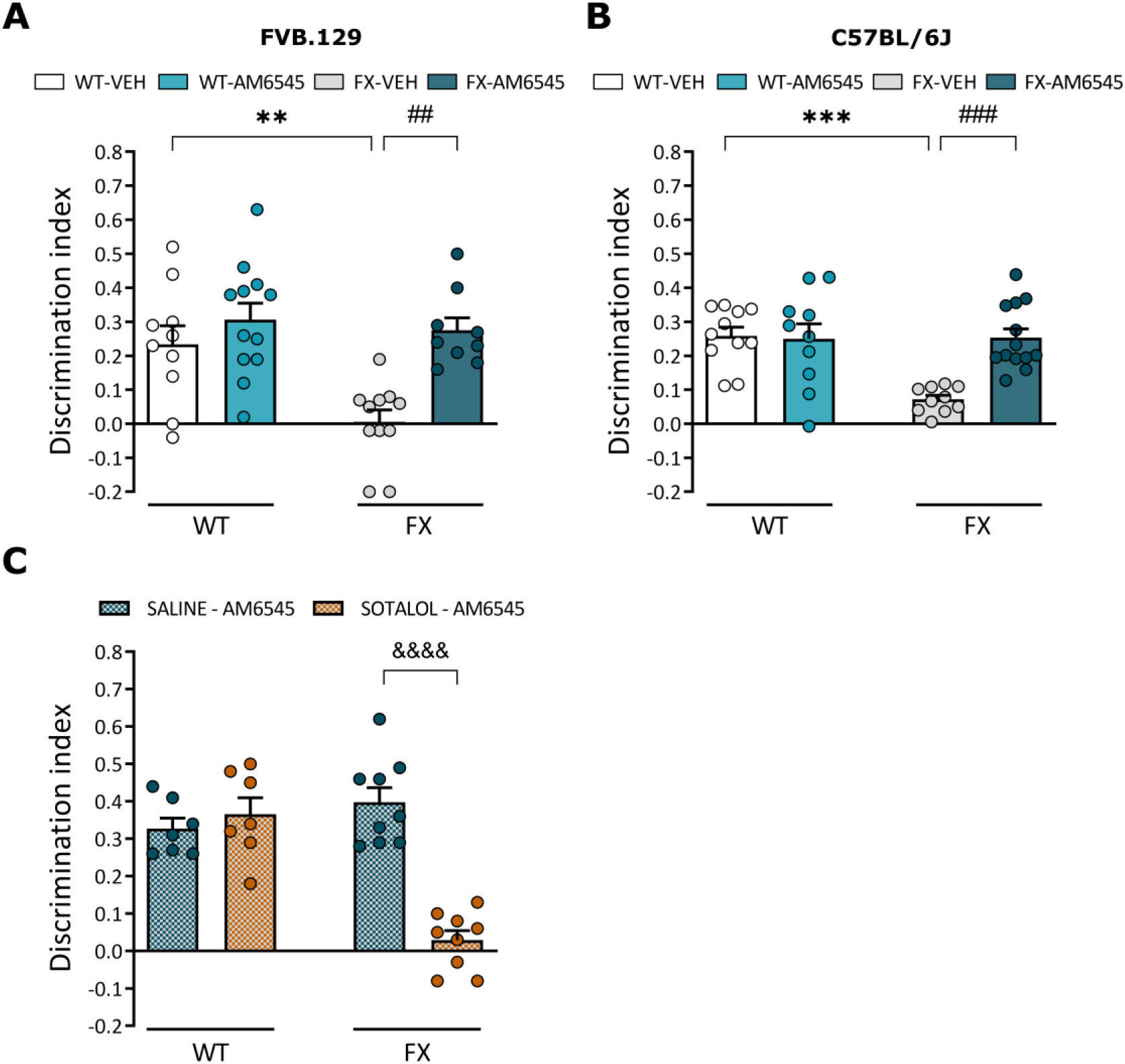
Memory rescue in FX mice by AM6545 treatment is mediated by peripheral noradrenergic signaling. **(A, B)** Discrimination index of WT and FX mice after sub-chronic administration of vehicle (VEH) or AM6545 (1 mg/kg, 7d) in **(A)** FVB.129 (*n* = 9-12) and **(B)** C57BL/6J (*n* = 10-13) genetic background. **(C)** Discrimination index of FVB.129 WT and FX mice pre-treated with saline or sotalol (10 mg/kg) before acute AM6545 (1 mg/kg) treatment (*n* = 7-9). Data are expressed as mean ± s.e.m. ** *p* < 0.01, *** *p* < 0.001 (genotype effect); ## *p* < 0.01, ### *p* < 0.001 (treatment effect); &&&& *p* < 0.0001 (pre-treatment effect) by two-way ANOVA followed by Bonferroni *post hoc*.

Since we previously demonstrated that AM6545 improves memory persistence through a peripheral adrenergic mechanism in naïve mice [13], we explored whether such a mechanism could also be involved in FX mice. We found that the acute mnemonic effect of AM6545 was inhibited by acute pre-treatment with the peripherally-restricted β-adrenergic antagonist sotalol (10 mg/kg) (two-way ANOVA, interaction: F(1,28) = 34.05, *p* = 0.000003; *post hoc* Bonferroni, FX-saline-AM6545 *vs.* FX-sotalol-AM6545 *p* < 0.0001) (Fig. 1C), pointing to a peripheral adrenergic mechanism in the pro-cognitive effect of AM6545 in FX mice. No differences in total exploration time were observed between pharmacological treatments or genotypes (Supplementary Fig. 2) discarding confounding effects on exploratory behavior. Thus, sub-chronic AM6545 treatment prevents the cognitive deficits observed in FX mice through a peripheral noradrenergic mechanism.

### Sub-chronic AM6545 treatment restores cellular alterations in FX hippocampus

The functional and structural synaptic alterations in FX mice that contribute to impairments in synaptic plasticity include aberrant mGluR5-LTD [5] and alterations in dendritic spines [27]. First, to test whether AM6545 treatment would restore the elevated hippocampal mGluR5-LTD, WT and FX mice were treated with AM6545 (1 mg/kg, 7d) or vehicle. The last AM6545 administration was performed 24h before brain extraction and hippocampal slice preparation for electrophysiological recording. After 10 min of baseline recording, bath application of DHPG (100 μM, 10 min) induced a transient acute depression, which is considered an electrophysiological readout of group I mGluR activation, followed by a small LTD in WT mice (Fig. 2A). However, we observed that the LTD induced by DHPG was significantly enhanced in vehicle-treated FX mice, which is consistent with previous reports on functional alterations in FXS [28]. Interestingly, we found that sub-chronic AM6545 treatment significantly ameliorated the enhanced mGluR5-LTD in hippocampal slices of FX mice (two-way ANOVA, interaction: F(1,23) = 4.769 *p* = 0.043; *post hoc* Bonferroni, WT-VEH *vs.* FX-VEH *p* = 0.002; FX-VEH *vs.* FX-AM6545 *p* = 0.003) (Fig. 2A-B).

**Figure 2:**
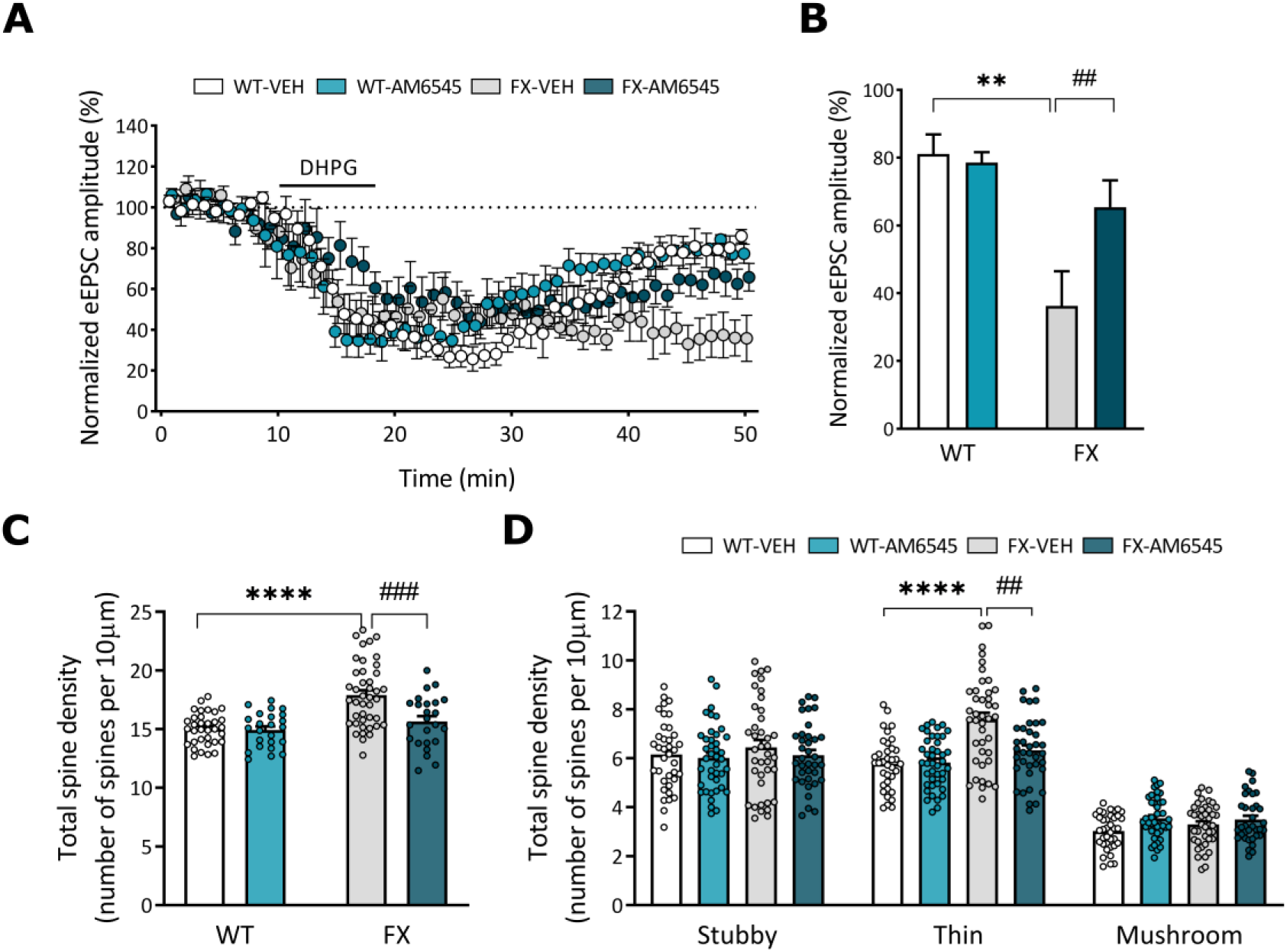
Sub-chronic AM6545 treatment ameliorates aberrant mGluR5-dependent LTD and normalizes dendritic spines alterations in FX mice. **(A)** Average time courses of the change in the amplitude of evoked excitatory postsynaptic currents (eEPSC) in hippocampal slices from FVB.129 mice treated for 7d with vehicle (VEH) or AM6545 (1 mg/kg, 7d) after bath application of DHPG (*n* = 6-7 mice). **(B)** Mean EPSC amplitude values of the last 5 min of recordings in **A**. **(C)** Quantification of the total number of spines in 10 μm of CA1 pyramidal dendrites of C57BL/6J WT and FX mice treated for 7d with VEH or AM6545 (*n* = 4-5 mice). **(D)** Morphological analysis of dendritic spines in the dendrites of pyramidal CA1 neurons in C57BL/6J WT and FX mice after pharmacological treatments (*n* = 4-5 mice). Each dot represents a hippocampal slice. Data are expressed as mean ± s.e.m. ** *p* < 0.01, **** *p* < 0.0001 (genotype effect); ## *p* < 0.01, ### *p* < 0.001 (treatment effect); by two-way ANOVA followed by Bonferroni *post hoc*.

Then, taking advantage of FX transgenic mice expressing the EGFP fluorescent protein under neuron-specific Thy1 promoter (Thy1-EGFP mice), we analyzed dendritic spine density and morphology of apical dendrites from CA1 Twenty-four hours after the last administration of sub-chronic AM6545 treatment. Vehicle-treated FX mice showed an increase in dendritic spine density compared to their WT littermates, that was normalized after sub-chronic AM6545 treatment (two-way ANOVA, interaction: F(1,120) = 7.728 *p* = 0.0063; *post hoc* Bonferroni, WT-VEH *vs.* FX-VEH *p* < 0.000001; FX-VEH *vs.* FX-AM6545 *p* = 0.00034) (Fig. 2C). Finally, we further classified the spines based on their morphology. We found that vehicle-treated FX mice presented an elevated density of thin (immature) spines that was restored after sub-chronic AM6545 treatment (two-way ANOVA, interaction: F(6,452) = 5.553 *p* = 0.000014; *post hoc* Bonferroni, WT-VEH *vs.* FX-VEH *p* < 0.0001; FX-VEH *vs.* FX-AM6545 *p* = 0.0010) (Fig. 2D). Therefore, sub-chronic AM6545 treatment normalizes the aberrant mGluR5-dependent LTD and dendritic spine density and morphology of CA1 pyramidal neurons in FX mice.

### Fecal microbiome of FX mice is sensitive to AM6545 treatment

To explore whether AM6545 could modify the gut microbiome, we investigated fecal microbiota in FX mice, and whether sub-chronic AM6545 treatment would modify fecal microbiota composition. To this end, we performed *16S* rRNA gene amplicon sequencing of feces samples in the same cohort of mice before and after vehicle or AM6545 administration, and analyzed the changes related either to the genotype or the treatment by paired comparisons (Supplementary Table 2). In terms of structural differences, no major changes were observed in α-diversity between the groups analyzed (Supplementary Fig. 3).

We observed differences in fecal microbiome composition between WT and FX mice in basal conditions, such as the reduction in *Lactobacillus* genus (Fig. 3A, Supplementary Table 3). Then, after behavioral experiments and sub-chronic vehicle administration, we found more changes between genotypes than in basal conditions. These changes included the reduction in the relative abundance of *Akkermansia* genus in vehicle-treated FX mice (Fig. 3B, Supplementary Table 4) compared to vehicle-treated WT mice.

**Figure 3:**
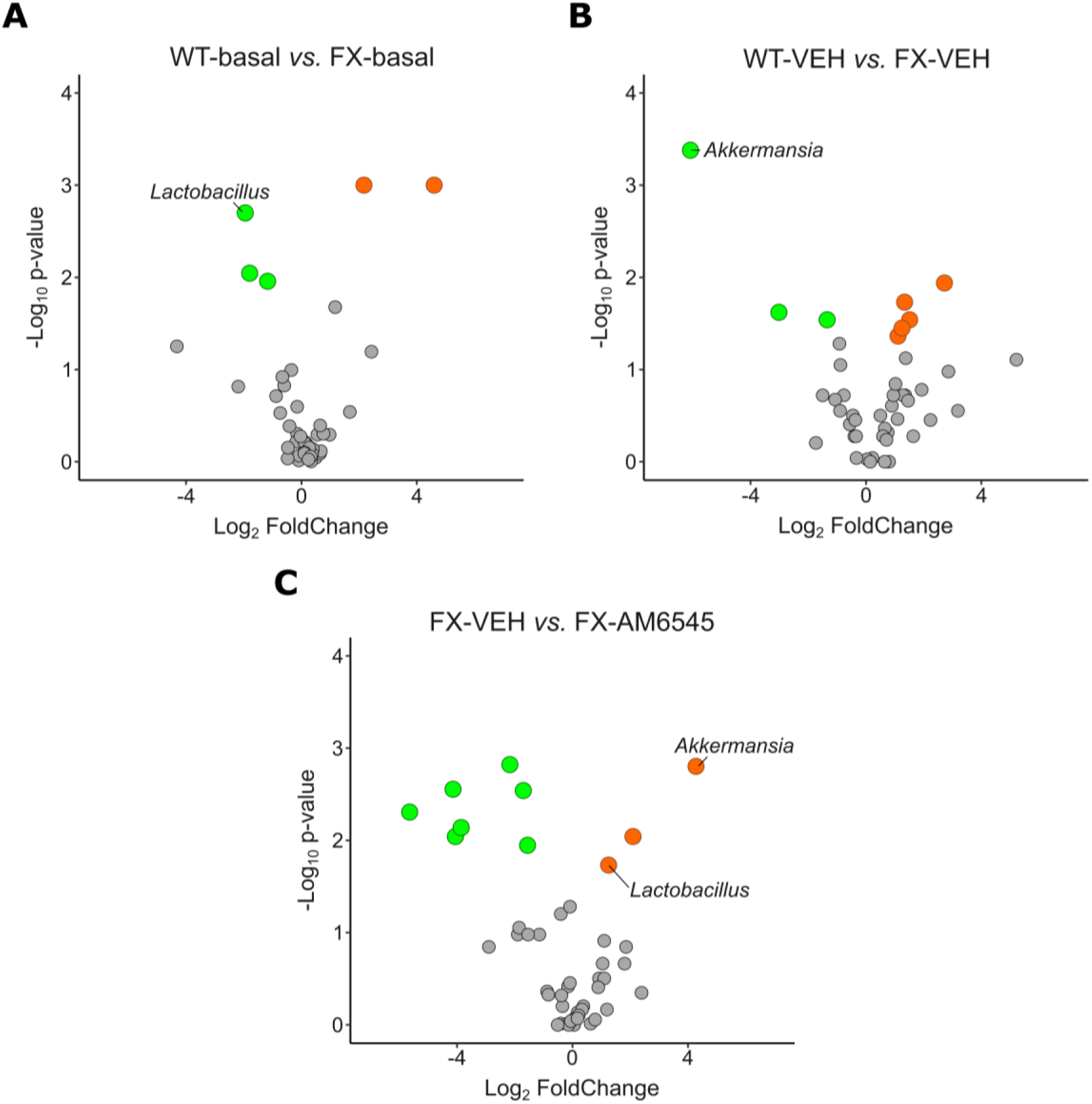
Fecal microbiome changes in FX mice after sub-chronic AM6545 treatment. Volcano plot showing differential abundance of microbiome composition in **(A)** C57BL/6J FX non-manipulated mice compared to WT non-manipulated mice (WT-basal *vs.* FX-basal), **(B)** C57BL/6J vehicle-treated FX mice compared to vehicle-treated WT mice (WT-VEH *vs.* FX-VEH) and **(C)** C57BL/6J FX mice sub-chronically treated with AM6545 (1mg/kg, 7d) compared to vehicle-treated FX mice (FX-VEH *vs.* FX-AM6545). Orange and green dots indicate enriched or depleted species in each comparison in terms of relative abundance at *genus* level. (WT-basal, *n* = 19; FX-basal, *n* = 20; WT-VEH, *n* = 10; FX-VEH, *n* = 10; FX-AM6545, *n* = 10). Significance is set at *p* < 0.05

Interestingly, sub-chronic AM6545 treatment results in an increase in bacteria belonging to both *Akkermansia* and *Lactobacillus* genera (Fig. 3C, Supplementary Table 5); pointing to the ability of AM6545 treatment to restore microbiota alterations observed in FX mice.

### Serotonin receptor 4 gene is upregulated in FX hippocampus after sub-chronic AM6545 treatment

To explore more in depth the synaptic impact of AM6545 in the context of FXS, we performed mRNA transcriptome analysis of hippocampal synaptoneurosomes. We treated FX mice for 7d with AM6545 or vehicle and obtained hippocampal samples 24h after the last AM6545 administration, just after the test phase of the NORT. DGE analysis revealed 7 genes significantly upregulated in FX mice treated with AM6545, whereas only *Dcp2*, was downregulated (Fig. 4A, Supplementary Table 6).

**Figure 4:**
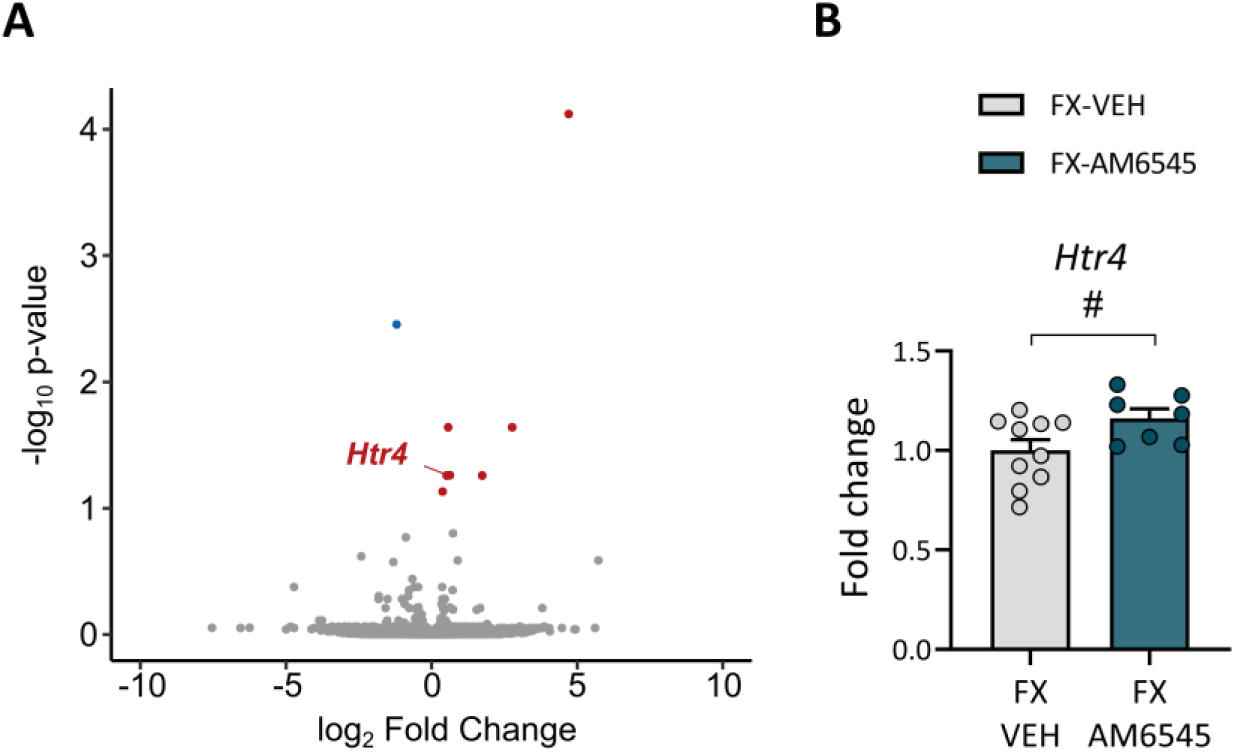
Sub-chronic AM6545 treatment upregulates serotonin 4 receptor in the hippocampus of FX mice. **(A)** Volcano plot showing differentially expressed genes in C57BL/6J FX mice treated with vehicle (VEH) or AM6545 (1 mg/kg, 7d). Red and blue dots represent significantly up or downregulated genes, respectively. (FX-VEH, *n* = 5; FX-AM6545, *n* = 4). Cut-off was set at |log2FC| > 0.2 and adjusted *p-value* < 0.1. **(B)** Synaptic hippocampal mRNA levels of *Htr4* in C57BL/6J FX mice treated for 7d with vehicle (VEH) or AM6545 (1 mg/kg, 7d) (*n* = 7-10). Data are expressed as mean ± s.e.m. # *p* < 0.05 (treatment effect) by Student’s t-test.

Among the upregulated genes, we focused on *Htr4*, since *Htr4*/5-HT4R expression/function is associated with hippocampal plasticity and processing of information [29]. We used an independent cohort of FX mice sub-chronically treated with AM6545 or vehicle, to validate the *in silico* result. We found, using qRT-PCR, that *Htr4* was upregulated in hippocampal synaptoneurosomes from AM6545-treated FX mice (Student’s t-test: *p* = 0.048) (Fig. 4B).

Interestingly, we found a positive correlation between discrimination index and *Htr4* levels from both RNA-sequencing (Pearson’s correlation coefficient = 0.858, *p* = 0.003) (Supplementary Fig. 4A) and qRT-PCR data (Pearson’s correlation coefficient = 0.485, *p* = 0.048) (Supplementary Fig. 4B). This indicates that serotonin receptor 4 mRNA levels increase in FX mice after AM6545 administration in a way that relates to the performance in NORT.

### Cognitive rescue of AM6545 in FX mice is dependent of a serotonergic mechanism

Based on *Htr4* correlation with memory in hippocampus, we hypothesized a possible implication of 5-HT4R in the alleviation of the cognitive impairment in FXS. To assess whether sub-chronic AM6545 treatment improves memory with the participation of a serotonergic signaling mechanism, FX mice were treated during 7d with vehicle or AM6545 and on the last day of treatment, they were pre-treated with the 5-HT4R antagonist GR125487 (10 mg/kg). Thus, immediately after the familiarization phase in the NORT, FX mice received an acute administration of GR125487 or saline, 20 min before the last AM6545 administration. When the test phase was performed 24h later, we observed that the pre-treatment with GR125487 abolished the mnemonic effect of sub-chronic AM6545 treatment in FX mice (two-way ANOVA, interaction: F(1,34) = 6.221, *p* = 0.018; *post hoc* Bonferroni, FX-saline-VEH *vs.* FX-saline-AM6545 *p* = 0.013; FX-saline-AM6545 *vs.* FX-GR125487-AM6545 *p* = 0.037) (Fig. 5). This effect was not related to modifications in the exploratory behavior since total exploration time did not significantly change (Supplementary Fig. 5). Together this indicates that the effect that AM6545 has on FX mice memory performance may involve 5-HT4R modulation.

**Figure 5:**
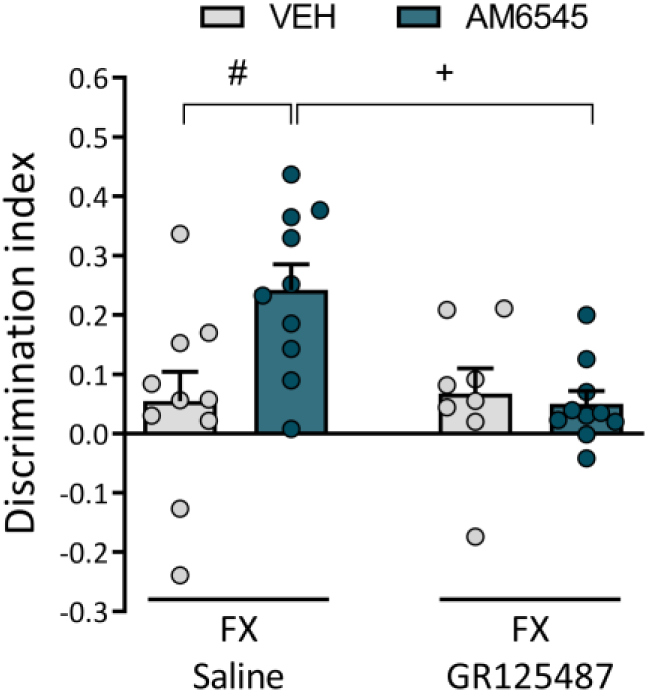
A serotonergic mechanism mediates the cognitive improvement of FX mice after sub-chronic AM6545 treatment. Discrimination index of C57BL/6J FX mice after sub-chronic vehicle (VEH) or AM6545 (1 mg/kg, 7d) administration and pre-treatment with saline or GR125487 (10 mg/kg) after the familiarization phase of the NORT (*n* = 8-10). Data are expressed as mean ± s.e.m. + *p* < 0.05 (pre-treatment effect); # *p* < 0.05 (treatment effect) by two-way ANOVA followed by Bonferroni *post hoc*.

## Discussion

In the present study we described that AM6545, a CB1R antagonist largely restricted to the periphery, rescues memory impairment, synaptic abnormalities and gut microbiota composition in FX mice. Moreover, we decipher that the memory rescue depends on peripheral β-adrenergic and 5-HT4R signaling in this FXS mouse model.

Previous research in our group has pointed to the endocannabinoid system as a potential therapeutic target in the treatment of FXS-associated cognitive deficits. Specifically, low doses of the systemic CB1R antagonist rimonabant, and the neutral CB1R antagonist NESS0327 showed improvements in several traits of FX mice such as the memory impairment, the elevated mGluR5-LTD, the alterations in dendritic spines or the susceptibility to audiogenic epileptic seizures [7,11]. Here, we aimed to target peripheral CB1R in the FXS mouse model, as we previously reported that the memory-enhancing effects of systemic CB1R inhibition with rimonabant in naïve mice can be reproduced by peripheral CB1R antagonists such as AM6545 [13,14].

First, we assessed memory performance in the NORT in mice that were sub-chronically treated with AM6545. Under these conditions, we found a significant amelioration of the memory deficits after the 7d treatment in two different genetic backgrounds, one outbred (FVB.129) and another inbred (C57BL/6J). This finding indicates a genuine overall effect of AM6545 that is not specific of the genetic background similarly as it was described for two different strains in naïve mice [14].

To further characterize the mnemonic effect of AM6545, we addressed whether peripheral β-adrenergic receptors could be involved, as it happens in naïve mice [13]. Acute pre-treatment with the β-adrenergic peripheral antagonist sotalol bluntly prevented the memory improvement of the acute peripheral treatment with AM6545 in FX mice.

Thus, sotalol effect in FX mice after AM6545 administration is reminiscent of its effect in preventing the memory improvement with AM6545 in naïve mice [13,14]. Since this goes in accordance with the modulation of cognitive performance by the activation of peripheral β-adrenergic receptors and/or afferent fibers from the vagus nerve [30,31], AM6545 effect in FX mice may also imply the participation of these fibers as in naïve mice [13]. Notably, vagus nerve electrostimulation increases noradrenaline levels in the hippocampus, and a reduction in noradrenergic transmission blocks the effects of vagus nerve stimulation in the central nervous system [32,33]. Moreover, our group also reported that auricular transcutaneous vagus nerve electrostimulation enhanced memory performance in FX mice [34], which reinforces the implication of vagal fibers afferents in memory improvement in FXS.

Interestingly, AM6545 normalizes the enhanced hippocampal mGluR5-LTD of FX mice and also restores the increased density of immature spines in the hippocampus. These cellular alterations have been well-described in FX mice [5,27] and have been typically associated to cognitive impairment in FXS [35]. The changes in synaptic function and structure of AM6545 are somehow reminiscent to the effect of sub-chronic treatment with rimonabant in which mGluR5-LTD and the number of immature spines in the hippocampus of FX mice were normalized [7,11]. Overall, these effects on synaptic plasticity in FX mice are consistent with the unanticipated hypothesis that rimonabant systemic effects can also be obtained targeting peripheral CB1R.

Given that AM6545 is a largely peripheral antagonist [36] and that alterations in gut microbiota have been described in FX mice [9], we next analyzed fecal microbiome composition. Here we reported a significant decrease in *Lactobacillus* and *Akkermansia* population in feces from FX mice compared to WT littermates. Notably, we also found that both reductions in abundance were restored after AM6545 treatment. Both genera have been recently pinpointed by others in a similar context. Treatment with *Lactobacillus* in mouse models of ASD restored social cognitive deficits and synaptic functions [37,38]. Indeed, a clinical trial is ongoing to evaluate the beneficial effects of increasing *Lactobacillus* species in children with FXS (NCT06279858). Moreover, it is described the implication of the vagus nerve in the behavioral and molecular changes induced by *Lactobacillus* in naïve mice [39], which agrees with the proposed mechanism of action of AM6545 [13]. Consistent with our results, *Akkermansia* levels were described to be low in FX mice [9] and when their levels increase, cognitive deficits are restored [40,41]. Furthermore, treatment with *Akkermansia muciniphila* in a nonalcoholic steatohepatitis rat model reverse cognitive impairment [42]. Although further research is needed to elucidate the role of these genera in FXS, some of the effects of AM6545 administration could be attained through the microbiota-gut-brain axis [43].

Our transcriptomic analysis of hippocampal synaptoneurosomes revealed few changes in gene expression. *Dcp2* downregulation, could indicate a higher concentration of this mRNA decapping protein in P-bodies [44], facilitating its interaction with excessive mRNAs found in FX synapses. Interestingly, we found the upregulation of *Htr4* in FX mice that received AM6545, strongly indicating a possible role of 5-HT4R in AM6545-mediated hippocampal changes. Importantly, 5-HT4R plays a relevant role in hippocampus-dependent learning and memory and the activation of these receptors has been related with cognitive enhancement [29]. The administration of 5-HT4R agonists enhance memory in naïve mice [45,46] and ameliorate cognitive deficits in mousemodels od Alzheimeŕs disease [47,48], while 5-HT4R decreased density has been associated to cognitive impairments [49,50]. In this regard, this receptor subclass has been suggested as neuronal marker for learning and memory [51].

Moreover, dysregulation in serotonin modulation has also been reported in ASD and FXS [52,53]. Recent studies reported a reduced expression of 5-HT1_B_R in subjects with FXS [54] and lower binding activity of 5HT1_A_R in FX mice [55]. Interestingly, the modulation of specific serotonin receptors (5-HT1R, 5-HT2R, 5-HT5R and 5-HT7R) have been described to ameliorate behavior and synaptic plasticity in FXS models [56–60]. However, changes in 5-HT4R have not been previously associated with FXS.

Finally, to further understand the role of 5-HT4R in AM6545 effect in FXS, we addressed whether the activation of 5-HT4R may interfere with memory performance in FX mice. Acute pre-treatment with the 5-HT4R antagonist GR125487, abolished the pro-cognitive effect of the sub-chronic AM6545 treatment in FX mice. In this regard, previous studies demonstrated a significant interaction between serotonergic and endocannabinoid systems [61]. Genetic or pharmacological CB1R inhibition has been shown to enhance serotonin levels and control the function and expression of serotonin receptors [62–64]. Specifically, low doses of 5-HT4R agonists reverse CB1R-induced memory impairment [65]. Moreover, activation of 5-HT4Rs has been described to increase intracellular cAMP levels, induce LTP and regulate neurotransmitter release in hippocampus [66,67]; which may explain why its activation is needed for the mnemonic effect of AM6545 in FX mice.

Altogether our results suggest that the pharmacological inhibition of CB1R with the largely peripheral compound AM6545 alleviates significant central and peripheral alterations in the FX mouse and pinpoint to the 5-HT4R as a central effector in the amelioration of cognitive dysfunction.

## Acknowledgements

We are grateful to Raquel Martín, Marta Linares, Dulce Real and Francisco Porrón for expert technical assistance and Lorena Galera-López and Laura Ciaran-Alfano for helpful discussion.

## Author contribution

L.R.-R., Behavioral and biochemical experiments, transcriptomic study, statistical analysis, graphs and writing of the manuscript; A.B.-M., Behavioral and biochemical experiments, statistical analysis and writing of the manuscript; S.M.-T., Behavioral experiments. I.M.-G., P.L.-R. and A.R.-M., Electrophysiological experiments; S.A. and M.G., Microbiota experiments. R.M., Supervision and funding of the study. A.O., Conceptualization, supervision, funding of the study and writing of the manuscript. All authors revised the final version of the manuscript.

## Data availability

The behavioral and biochemical data that support the findings of this study are available from the corresponding author upon reasonable request. Transcriptomic data in the form of raw fastq files have been deposited on GEO database (GSE278037).

