## Supplementary Information for "A serotonergic mechanism is involved in the pro-cognitive effect of AM6545 treatment in a mouse model of fragile X syndrome"

### Supplementary Material

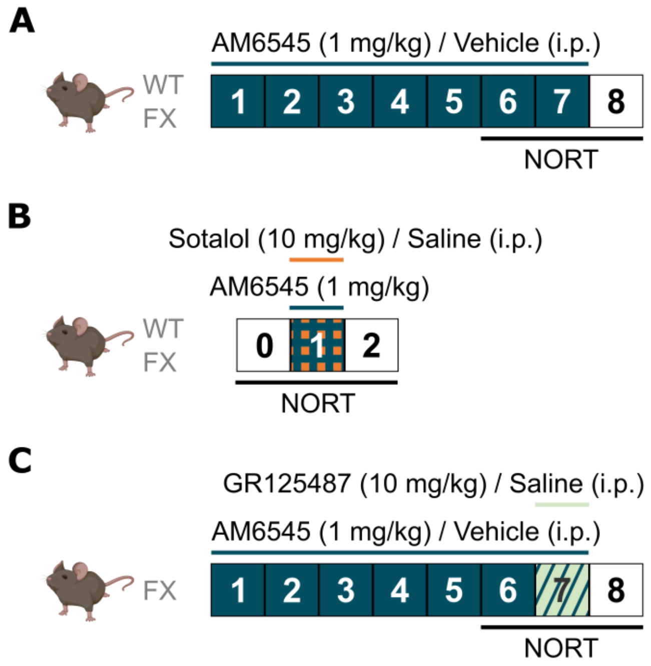


**Supplementary Figure 1: Schedule of administration of pharmacological approaches for behavioral tests. (A)** Sub-chronic AM6545 (1 mg/kg, 7d) or vehicle administration to WT and FX mice. **(B)** Acute AM6545 administration after acute pre-treatment with sotalol (10 mg/kg) or saline to WT and FX mice. **(C)** Sub-chronic AM6545 or vehicle administration combined with acute pre-treatment with GR125487 (10 mg/kg) or saline administration to FX mice. Samples were collected 24 h after the last AM6545 administration of the sub-chronic treatment (day 8 on the scheme).

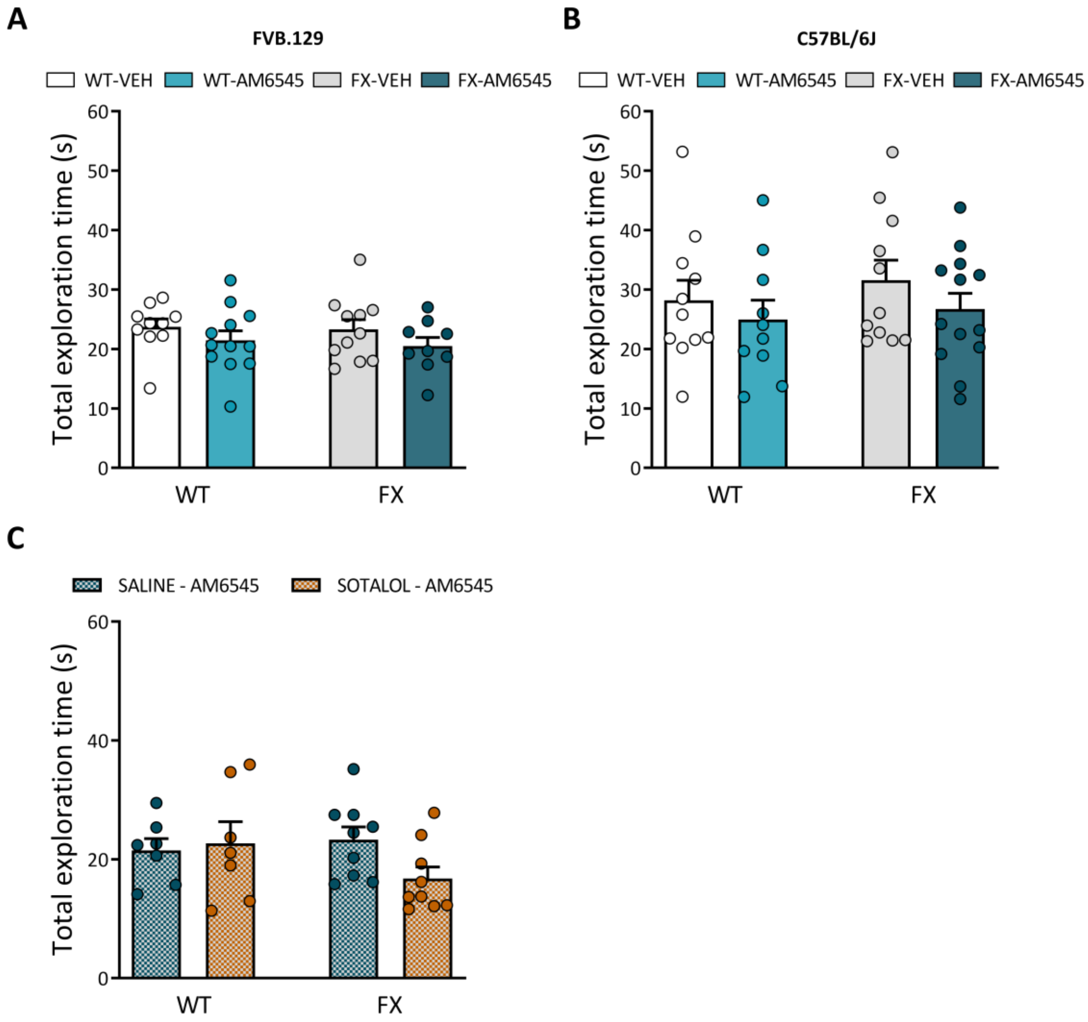


**Supplementary Figure 2: Pharmacological interventions do not modify total exploration time in NORT. (A, B)** Total exploration time of WT and FX mice after sub-chronic administration of vehicle (VEH) or AM6545 (1 mg/kg, 7d) in **(A)** FVB.129 (*n* = 9-12) and **(B)** C57BL/6J (*n* = 10-13) genetic background. **(C)** Total exploration time of FVB.129 WT and FX mice pre-treated with saline or sotalol (10 mg/kg) before acute AM6545 (1 mg/kg) treatment (*n* = 7-9). Data are expressed as mean ± s.e.m. Statistical significance was calculated by two-way ANOVA.

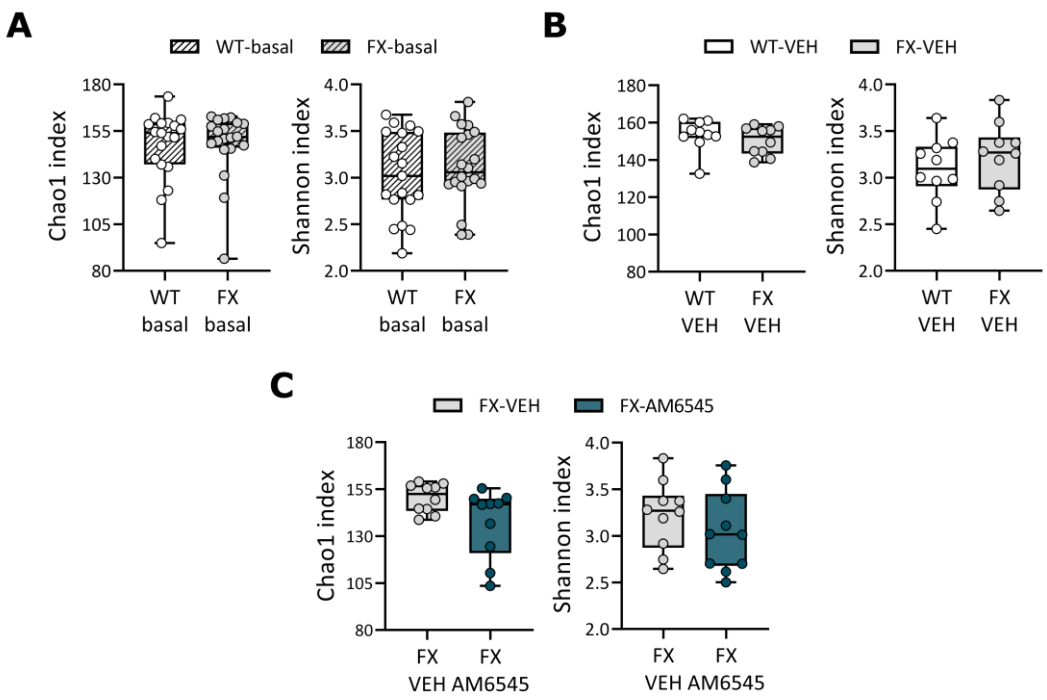


**Supplementary Figure 3: Diversity is not modified in C57BL/6J FX mice neither in basal, nor after sub-chronic AM6545 treatment. (A-C)** Chao1 and Shannon indexes of microbiome from **(A)** non-manipulated WT and FX mice (basal), **(B)** WT and FX mice after vehicle (VEH) administration and **(C)** FX mice after sub-chronic vehicle or AM6545 (1 mg/kg, 7d) administration. (WT-basal, $n$ = 19; FX-basal, $n$ = 20; WT-VEH, $n$ = 10; FX-VEH, $n$ = 10; FX-AM6545, $n$ = 10). Statistical significance was calculated by two-tailed Student's t-test.

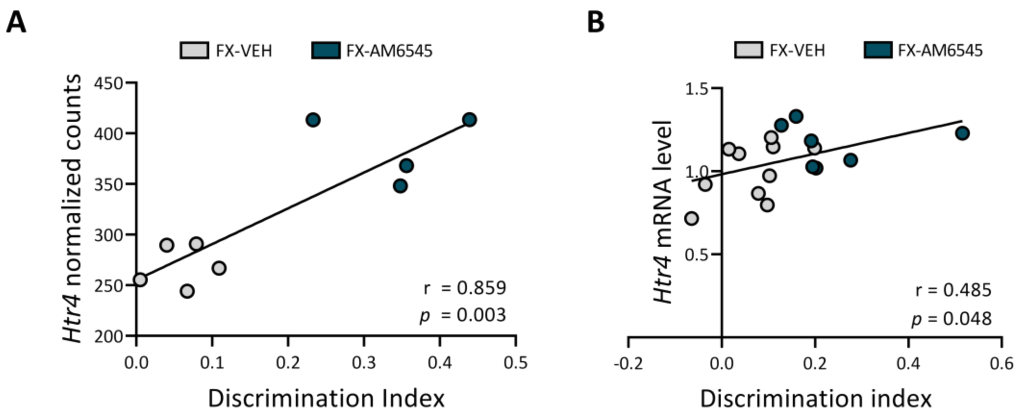


**Supplementary Figure 4: *Htr4* levels correlate with memory performance in the NORT.** Correlations of discrimination index of C57BL/6J FX mice after vehicle (VEH) or AM6545 (1 mg/kg, 7d) administration and hippocampal synaptoneurosomes *Htr4* relative expression from **(A)** RNA-sequencing and from **(B)** qRT-PCR validation.

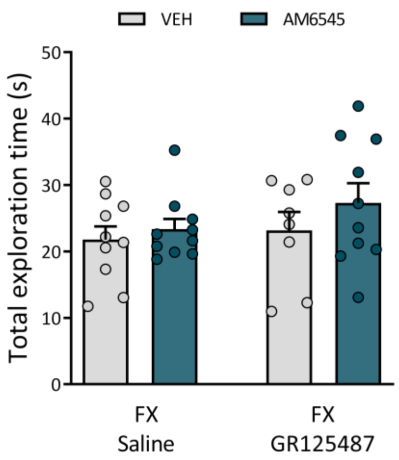


**Supplementary Figure 5: GR125487 does not modify total exploration time in NORT.** Total exploration time of C57BL/6J FX mice after sub-chronic vehicle (VEH) or AM6545 (1 mg/kg, 7d) administration and pre-treatment with saline or GR125487 (10 mg/kg) after the familiarization phase of the NORT (*n* = 8-10). Data are expressed as mean ± s.e.m. Statistical significance was calculated by two-way ANOVA.

***NeuronStudio parameters***

| ***Spine type*** | ***Head/neck ratio*** | ***Length/head ratio*** | ***Head size*** |
|---|---|---|---|
| Mushroom | > 1.1 | -- | > 0.350 µm |
| Stubby | < 1.1 | < 2.5 | -- |
| Thin | -- | > 2.5 | < 0.350 µm |

**Supplementary Table 1**. **NeuronStudio parameters for spines morphologic classification**. For each spine, NeuronStudio calculated head/neck ratio, length/head ratio and head size and then, when following both criteria for these parameters, spines were classified in mushroom, stubby and thin.

***Experimental groups***

| ***Group name*** | ***Manipulation*** | ***Comparison*** | ***Experiment*** | ***Group size*** |
|---|---|---|---|---|
| Non-manipulated (basal) | Non-manipulated WT and FX mice | WT-basal *vs.* FX-basal | Fecal microbiota | WT-basal, n = 19<br>FX-basal, n = 20 |
| Genotype effect | WT and FX mice after behavior and 7d vehicle administration | WT-VEH *vs.* FX-VEH | Fecal microbiota | WT-VEH, n = 10<br>FX-VEH, n = 10 |
| Treatment effect in FX | FX mice after behavior and 7d vehicle or AM6545 (1 mg/kg) administration | FX-VEH *vs.* FX-AM6545 | Fecal microbiota | FX-VEH, n = 10<br>FX-AM6545, n = 10 |
| Treatment effect in FX | FX mice after behavior and 7d vehicle or AM6545 (1 mg/kg) administration | FX-VEH *vs.* FX-AM6545 | Synaptic gene expression | FX-VEH, n = 5<br>FX-AM6545, n = 4 |

**Supplementary Table 2**. **Overview of the experimental groups**. Experimental design and group comparisons for fecal microbiota and differential gene expression analysis.

**WT-basal *vs.* FX-basal**

| ***Genus*** | ***Log₂FC*** | ***P-values*** | ***Genus*** | ***Log₂FC*** | ***P-values*** |
|---|---|---|---|---|---|
| *Lactobacillus* | -1.94 | 2.00E-03 | *Pseudomonas* | 4.60 | 1.00E-03 |
| *Lachnospiraceae_UCG_006* | -1.79 | 9.00E-03 | *Colidextribacter* | 2.16 | 1.00E-03 |
| *Enterorhabdus* | -1.17 | 1.10E-02 | | | |

**Supplementary Table 3**. **Differential abundance in fecal samples from non-manipulated WT and FX mice**. Summary of differential abundance in fecal samples from FX mice compared to WT mice, both non-manipulated. (WT-basal, n = 19; FX-basal, n = 20). Statistical significance was calculated through Mann-Whitney U test and taxonomic profiles at genus level were presented with the cut-off settled at p-value<0.05 and $|log_2FC| > 1$.

**WT-VEH *vs.* FX-VEH**

| *Genus* | *$Log_2FC$* | *P-values* | *Genus* | *$Log_2FC$* | *P-values* |
|---|---|---|---|---|---|
| *Akkermansia* | -6.08 | 4.18E-04 | *Mycoplasma* | 2.72 | 1.15E-02 |
| *Lachnospiraceae_UCG_008* | -3.02 | 2.39E-02 | *Helicobacter* | 1.34 | 1.85E-02 |
| *Roseburia* | -1.35 | 2.88E-02 | *Anaerotruncus* | 1.51 | 2.88E-02 |
| | | | *ASF356* | 1.25 | 3.55E-02 |
| | | | *Enterorhabdus* | 1.11 | 4.33E-02 |

**Supplementary Table 4**. **Differential abundance in fecal samples from vehicle-treated WT and FX mice**. Summary of differential abundance in fecal samples from FX mice treated with vehicle compared to WT mice treated with vehicle. (WT-VEH, n = 10; FX-basal, n = 10). Statistical significance was calculated through Mann-Whitney U test and taxonomic profiles at genus level were presented with the cut-off settled at p-value<0.05 and $|log_2FC| > 1$.

**FX-VEH *vs.* FX-AM6545**

| *Genus* | *$Log_2FC$* | *P-values* | *Genus* | *$Log_2FC$* | *P-values* |
|---|---|---|---|---|---|
| *Helicobacter* | -2.17 | 1.50E-03 | *Akkermansia* | 4.28 | 1.57E-03 |
| *Pseudomonas* | -4.14 | 2.79E-03 | *Lachnospiraceae_UCG_006* | 2.10 | 9.08E-03 |
| *Anaerotruncus* | -1.71 | 2.88E-03 | *Lactobacillus* | 1.25 | 1.85E-02 |
| *Faecalibaculum* | -5.64 | 4.93E-03 | | | |
| *Mycoplasma* | -3.86 | 7.26E-03 | | | |
| *Mitochondria* | -4.06 | 9.08E-03 | | | |
| *Lachnospiraceae_UCG_001* | -1.56 | 1.13E-02 | | | |

**Supplementary Table 5**. **Differential abundance in fecal samples from FX mice treated with AM6545 or vehicle**. Summary of differential abundance in fecal samples from FX mice treated with AM6545 (1 mg/kg, 7d) compared to FX mice treated with vehicle. (FX-VEH, n = 10; FX-AM, n = 10). Statistical significance was calculated through Mann-Whitney U test and taxonomic profiles at genus level were presented with the cut-off settled at p-value<0.05 and $|log_2FC| > 1$.

**FX-VEH *vs.* FX-AM6545**

| *Gene name* | *$Log_2FC$* | *Adj. p-value* | *Gene name* | *$Log_2FC$* | *Adj. p-value* |
|---|---|---|---|---|---|
| *Dcp2* | -1.200 | 3.50E-03 | *Ttn* | 4.705 | 7.59E-05 |
| | | | *Topbp1* | 0.571 | 2.29E-02 |
| | | | *Fanca* | 2.766 | 2.29E-02 |
| | | | *Ctc1* | 0.628 | 5.48E-02 |
| | | | *Htr4* | 0.522 | 5.51E-02 |
| | | | *Gm14308* | 1.740 | 5.51E-02 |
| | | | *Lmo7* | 0.379 | 7.36E-02 |

**Supplementary Table 6**. **Differential gene expression in hippocampal synaptoneurosomes from FX mice treated with AM6545 or vehicle**. Summary of differentially expressed genes in hippocampal synaptoneurosomes from FX mice treated with AM6545 (1 mg/kg, 7d) compared to FX mice treated with vehicle. Threshold for significance was. (FX-VEH, n = 5; FX-AM6545, n = 4). Statistical significance was calculated by Benjamini-Hochberg adjustment following Wald test analysis with the cut-off settled at adjusted p-value < 0.1 and $|log_2FC| > 0.2$.
